# Human length telomeres restrict the regenerative potential of hematopoietic stem cells in mice

**DOI:** 10.1101/2025.08.26.672314

**Authors:** Melissa Rowe, Joanna Tober, Vivian Ortiz, Riham Smoom, Yehuda Tzfati, Nancy Speck, Klaus H. Kaestner

**Author notes:** These authors contributed equally (Melissa Rowe, Joanna Tober).

## Abstract

Extremely short telomeres cause bone marrow failure in telomere biology disorder (TBDs) patients. Here, we employed the recently developed ‘Telomouse’ with human-length telomeres resulting from a single amino acid substitution in the helicase Rtel1 (*Rtel1*^M492K/M492K^) to determine the effects of the short telomeres on the bone marrow and hematopoiesis. Under homeostatic conditions, Telomice have notably short telomeres but normal hematopoiesis. However, when forced to repopulate following repeated treatment with 5-fluoro-uracil or upon bone marrow transplantation into lethally irradiated mice, bone marrow progenitor cells are significantly depleted in Telomice compared to wild-type controls. This effect is associated with increased frequency of telomere repeat arrays too short to be detected by fluorescence in situ hybridization in the bone marrow of Telomice.

## Introduction

Telomeres are repeated sequences of DNA complexed with shelterin proteins at the ends of the chromosomes, which protect genome integrity and function (de Lange 2025). Telomere length is especially important in highly replicative tissues such as the bone marrow, where hematopoietic stem cells replicate every 25-50 weeks in humans (Catlin et al. 2011) and every 2.5 weeks in mice (Abkowitz et al. 2000). This self-renewal capability is dependent on the enzyme telomerase, which adds telomeric repeats to compensate for the shortening that occurs during each cycle of DNA replication. Human telomerase is active predominantly in germ, stem and progenitor cells. The importance of telomere length in highly regenerative tissues is emphasized in humans with genetic telomere biology disorders (TBDs) such as dyskeratosis congenita (DC) and Hoyeraal-Hreidarsson Syndrome (HHS). The predominant symptom and main cause of mortality in DC and HHS is bone marrow failure, underscoring the impact of dysfunctional telomeres on replicative cell niches such as the hematopoietic system (Glousker et al. 2015).

Unfortunately, human telomere biology is not well recapitulated by the common *M. musculus* lab strains, which have telomere repeat arrays that are about 5 times longer than those present in humans (Smoom et al. 2023). As a result, *M. musculus* cells can tolerate many cycles of replication even in the absence of telomerase activity before telomeres become critically short. Even mice null for the telomerase RNA component *mTR*, lacking all telomerase activity, are viable until the sixth generation of intercrossing (Blasco et al. 1997); in contrast, complete Telomerase deficiency is incompatible with life in humans (Armanios et al. 2005). In order to bridge this gap between human and mouse biology, we recently derived a new *M. musculus* C57BL/6 mouse strain with generationally stable human-length telomere repeat arrays, the ‘Telomouse’ (Smoom et al. 2023). This was accomplished by introducing a point mutation into the *Rtel1* gene that changes the methionine at position 492 of the RTEL1 helicase to a lysine (M492K). While these mice have telomeres typical of a healthy human, their length is well below what is normal for their species. Here, we tested the hypothesis that the shortened telomeres present in the Telomouse bone marrow might limit the regenerative capacity of the hematopoietic system.

## Results and Discussion

In order to evaluate if shortened telomeres limit the regenerative capacity in late generation Telomice (*Rtel1*^M492K/M492K^; between F11 and F17) compared to otherwise isogenic *M. musculus* C57BL/6 controls, we first determined bone marrow function at baseline under homeostatic conditions. We isolated bone marrow from young (2 months) and aged (12 months) mice and determined the proportion of various stem and progenitor lineages using flow cytometry (Fig. 1A, B; see Supplementary Figure 1 for flow cytometry gating strategy, and Supplementary Table 1 for a list of antibodies employed). We employed antibodies to the SLAM (signaling lymphocyte activation molecule) family proteins CD150 and CD48 first described by Kiel and colleagues for the definition of progenitor cell lineages (Kiel et al. 2005). While aged mice showed a relative increase in phenotypic long-term repopulating HSCs (LSK^150+48-^) cells and a relative decrease in lymphoid-biased progenitor (LSK^150-48+^) cells compared to young mice, there were no differences by genotype (Fig. 1A, B). Thus, under homeostatic conditions, the reduced telomere repeat length of Telomice does not impair bone marrow function.

**Figure 1.**
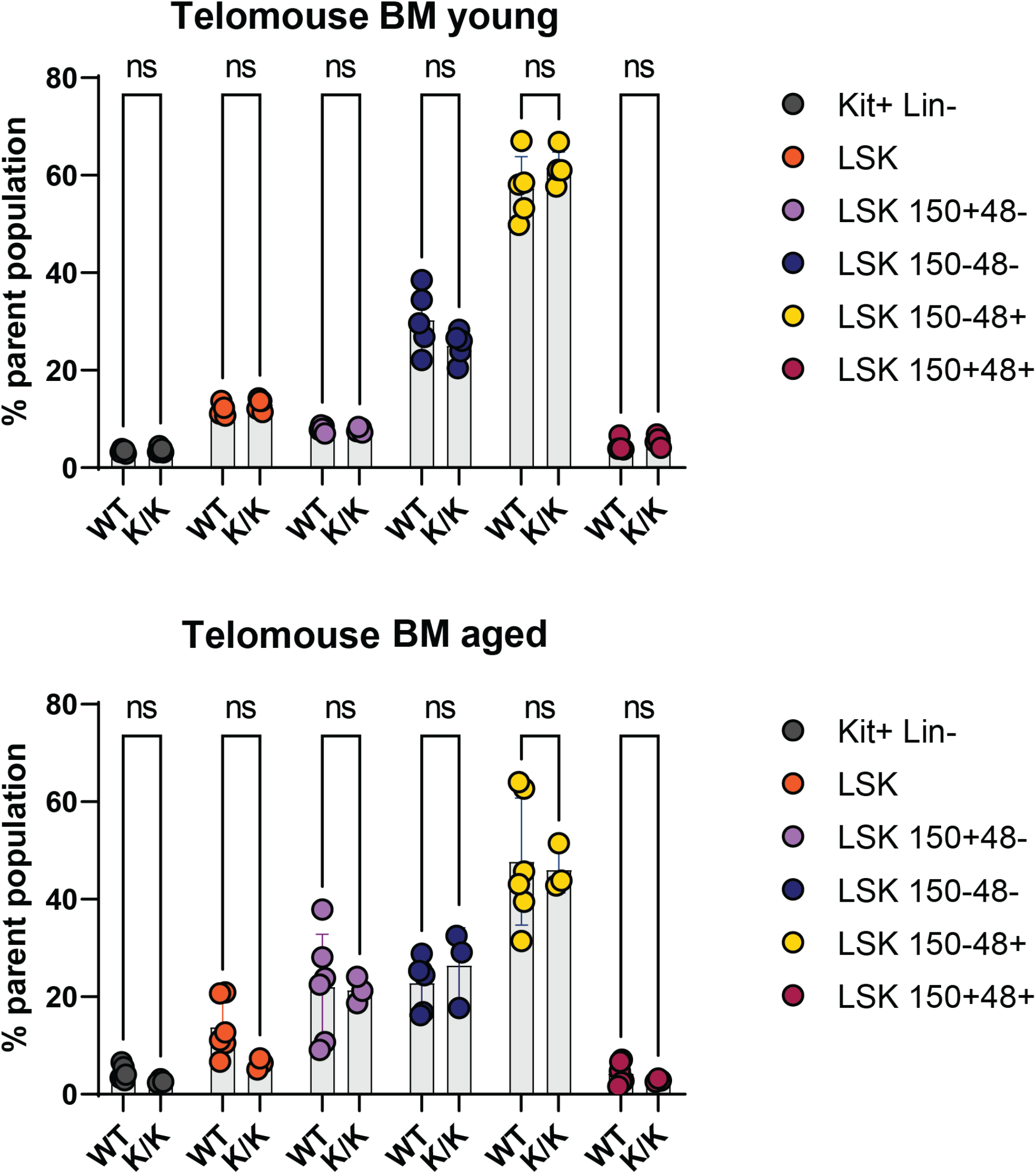
Homeostatic bone marrow function is normal in Telomice (K/K). The bone marrow composition of (A) young (2 months) and (B) aged mice (1 year) was determined using flow cytometry. Kit+ Lin-, lineage-negative bone marrow progenitors (lineage minus, c-Kit+ cells). Lineage markers = B220+, B-cells; CD3e+, T-cells; CD11b+; Macrophages/Granulocytes; Gr1, Granulocytes), (gray). LSK, hematopoietic stem and multipotent progenitor cells (lineage-negative, Sca-1+c-Kit+) (orange). Other stem/progenitor bone marrow populations are shown as percentages of the LSK population; LSK CD150+CD48-(purple): phenotypic long-term repopulating hematopoietic stem cells (HSC); LSK CD150-CD48-(blue): multipotent progenitors with transient multilineage reconstitution potential; LSK 150-48+ (yellow) and LSK CD150+CD48+ (red): lineage-restricted progenitors. Statistical analysis was performed using two-way ANOVA, multi-comparison Sidak’s test. For the young cohorts: 5 WT mice and 5 Telomice (F14) were assayed. For the aged cohort, 5 WT mice and 3 Telomice (F12) were analyzed.

Having established that there are no significant differences in bone marrow composition between Telomice and WT mice at baseline homeostatic conditions, we next sought to test the ability of bone marrow cells to regenerate under replicative stress. To this end, Telomice and WT mice were dosed with 5-Fluorouracil (5-FU), a cancer-treatment drug known to deplete bone marrow cells (Radley and Scurfield 1979). 5-FU was administered to groups of Telomice and WT mice on days 0, 7, and 14, and bone marrow collected from all mice on day 21. Peripheral blood was also harvested at 2–3-day intervals over the course of the entire experiment (Fig. 2A). Throughout the experimental timeline, some mice had to be euthanized for humane reasons. This occurred significantly more frequently in Telomice than in control mice (Fig. 2B). Notably, humanely euthanized mice had pale extremities and low bone marrow cell recovery upon harvesting (data not shown). Over the course of the experiment, the percentage of kit^+^/CD45^+^ progenitor cells increased in the peripheral blood in both groups, indicative of the 5-FU effect, with no significant differences between WT and Telomice (Fig. 2C; see Supplementary Figure 2 for flow cytometry gating strategy). However, when we analyzed the bone marrow on day 21 of the study, we noted significantly lower percentages of lineage-negative (Lin^-^), Sca1+ Kit^+^ (LSK) hematopoietic stem cells in Telomice compared to WT mice (Fig. 2D). Of these, in particular LSK CD150+ CD48-, i.e. long-term repopulating hematopoietic stem cells (Kiel et al. 2005), and transient multilineage progenitor LSK CD150-CD48-populations were reduced in frequency in the Telomice compared to WT mice (Fig. 2D). These findings indicate that the shortened telomere repeats present in Telomice limit the ability of bone marrow stem cells to respond to strong regenerative stress.

**Figure 2.**
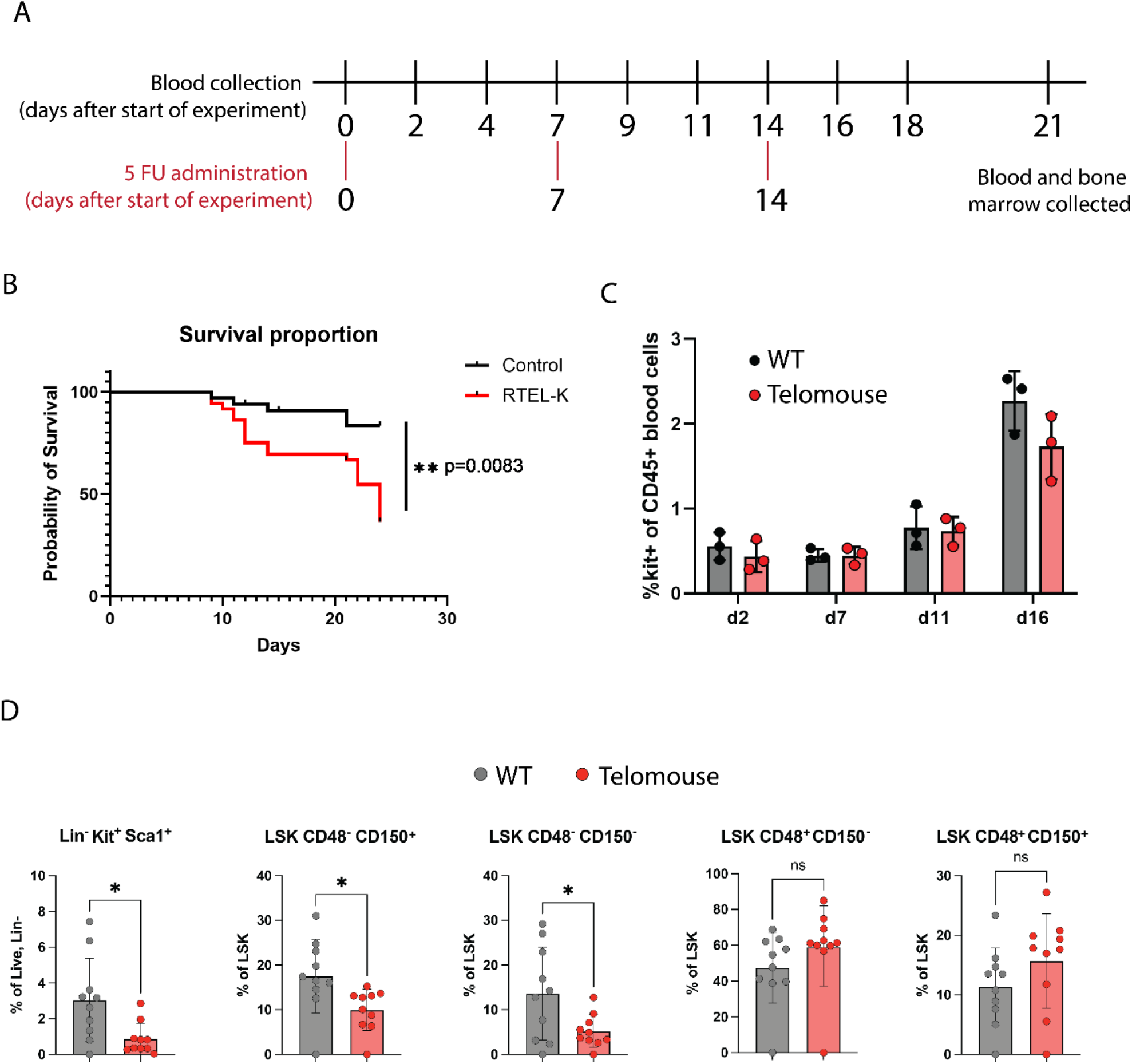
The bone marrow of Telomice does not recover effectively from 5-FU treatment. (A) Experimental timeline. Day 0 represents the collection of baseline peripheral blood and the first injection of 5-FU. Additional 5-FU injections occurred on days 7 and 14 (red lines). Peripheral blood collection from 2-3 mice of each genotype occurred on days indicated by vertical black lines. Peripheral blood and bone marrow were collected from all mice on day 21. (B) Survival curves for WT and Telomice. Significance was determined using the Mantel-Haenszel method. N= 34 WT mice and 36 Telomice (F15-17) (C) The percentage of CD45+ cells in peripheral blood which are also c-Kit+ in Telomice and WT mice over the course of the experiment. N= 10 WT mice and 10 Telomice (F13). (D) Comparisons of Telomouse and WT cell populations in the whole, non-lineage depleted, bone marrow at day 21. Student’s T-test was used for statistical analysis. Stem and progenitor cells were identified as in Fig. 1. N = 10 WT mice and 10 Telomice (F15).

To directly compare the regenerative ability of Telomouse bone marrow to that of wild type *Mus musculus* C57BL/6 mice, we performed a competitive bone marrow repopulation study (Fig. 3A). Bone marrow was harvested from 8–10-week-old CD45.2+ Telomice and 8–10-week-old CD45.1/CD45.2+ WT mice, and a mixture of one million cells each of the two populations was administered to lethally irradiated CD45.1+ hosts via retro-orbital injection. Peripheral blood and bone marrow were collected 16 weeks post-transplant. Cells were stained for common immune and hematopoietic stem cell markers and analyzed based on positivity for CD45.1 and CD45.2 to differentiate the donor contributions. In the peripheral blood, there was a slight reduction in the contribution of bone marrow from Telomouse donors to T-cells, macrophages, and total CD45+ cells compared to WT donors (Fig. 3B; see Supplementary Figure 3 for flow cytometry gating strategy). In the bone marrow, there was a higher contribution by Telomouse donors to CD45+ cells due to a large contribution to the granulocyte population (Fig. 3C). When analyzing the stem cell compartment, we noted nearly equal contribution of the two donor groups to the various stem and progenitor populations, though the Telomice provided significantly fewer LSK CD150+ CD48+ cells to the reconstituted bone marrow. LSK CD150+ CD48+ cells are considered to be lineage-restricted progenitors (Oguro et al. 2013). Overall, the bone marrow of young Telomice mice was not dramatically impaired in its ability to repopulate the bone marrow of lethally irradiated mice.

**Figure 3.**
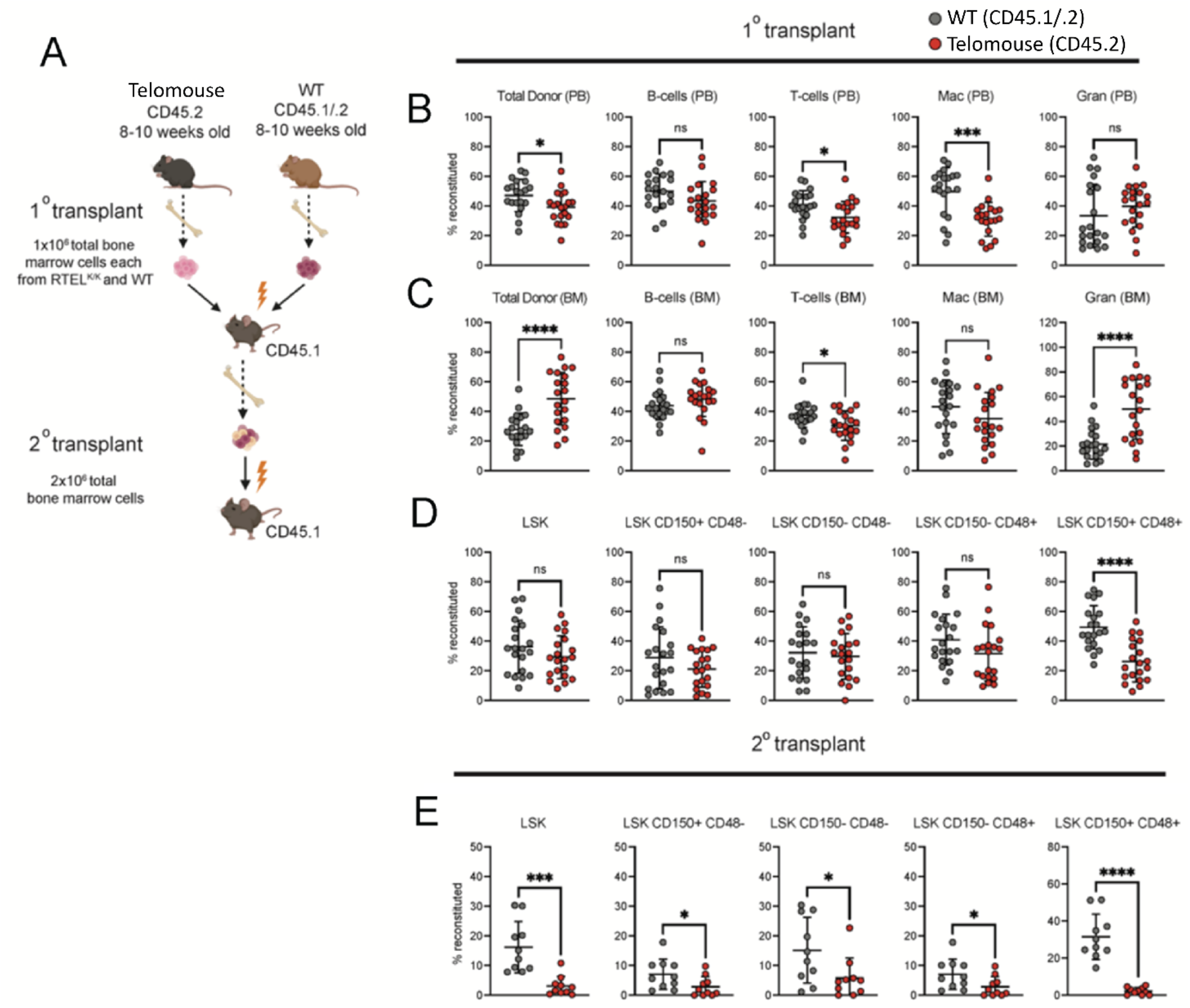
Young Telomice are competent to restore the hematopoietic system of a lethally irradiated recipient on initial but not serial transplantation. (A) Schema of experimental design. One million bone marrow (BM) cells each from Telomice (CD45.2^+^) and WT mice (CD45.1^+^/CD45.2^+^) were combined and transplanted into irradiated recipient mice. Sixteen weeks later, 2 x 10^6^ BM cells from 5 primary recipients were transplanted into 10 secondary recipients, (2 replicates for each primary recipient). (B) Telomice and WT donor contribution to peripheral blood (PB) cell lineages following 16 weeks of repopulation performed by gating donor cells (CD45.1^+^/CD45.2^+^ WT and CD45.2^+^ Telomice) within the total B-cell (CD19^+^CD3e^-^), T-cell (CD19^-^ CD3e^+^), macrophage (Mac, CD11b^+^Gr1^-^) and granulocyte (Gran, CD11b^+^Gr1^+^) populations (the total population also includes residual CD45.1+ recipient cells). (C) Telomice and WT donor contribution to B-cells, T-cells, macrophages, and granulocytes at 16 weeks of repopulation, determined as described in panel B. (D) Analysis of the hematopoietic stem cell compartment of recipient mice 16 weeks after bone marrow transplant. Stem and progenitor cells were identified as in Fig. 1. N= 20 WT mice and 20 Telomice (F11). (E) Contribution of WT and Telomouse donor cells to the hematopoietic stem and progenitor cell compartment of the bone marrow of recipient mice following secondary transplant. Percentages represent the proportions of WT-and Telomice-derived cells relative to all cells (CD45.2+, CD45.1+/CD45.2+, and CD45.1+). Student’s T-test was used for statical analysis. N = 5 WT and 5 Telomouse donors, 2 replicates of each donor (F11). ****, p ≤ 0.0001; ***, p ≤ 0.001; *, p < 0.05

Next, we sought to stress the regenerative potential of the Telomouse^-^derived hematopoietic stem cells even further by forcing the cells to undergo additional rounds of cell division. To this end, bone marrow from the original transplant recipient mice, containing marrow with both Telomouse CD45.2+ cells and WT CD45.1/CD45.2+ cells, was serially transplanted into a second irradiated CD45.1+ host mouse (Fig. 3A). Remarkably, stem and progenitor populations in the bone marrow of the secondary recipient exhibited dramatic reductions of the Telomouse contribution to the hematopoietic stem and multipotent progenitor cell pools (Fig. 3E). Thus, by forcing the hematopoietic stem cell compartment of Telomice to cycle multiple times, we uncovered a regenerative defect in these mice.

We sought to apply a second model to test if the regenerative capacity of the Telomouse bone marrow is limited by shortened telomeres. We repeated the competitive repopulation assay using bone marrow from aged mice (Fig. 4A). Bone marrow was harvested from one year old CD45.2+ WT or Telomice, and from 8–10-week-old CD45.1/CD45.2+ WT mice, and an equal mixture of the two combinations of young and old populations was administered to irradiated CD45.1+ hosts via retro-orbital injection. Peripheral blood and bone marrow were collected 16 weeks post-transplant. Cells were stained and sorted as described above Ammonium-Chloride-Potassium (Fig. 4B; see Supplementary Figure 3 for flow cytometry gating strategy). When comparing the regenerative functionality of the two aged bone marrow populations, there were significantly fewer phenotypic HSCs (LSK CD150+ CD48-) from aged Telomouse donors than from aged WT donors present in their respective hosts (Fig. 4B). We also analyzed lymphocytes, macrophages, and granulocytes in the peripheral blood from the competitive reconstitution study. Aged Telomouse donors were significantly less efficient in contributing to all hematopoietic lineages than aged wild type donors (Fig. 4C).

**Figure 4.**
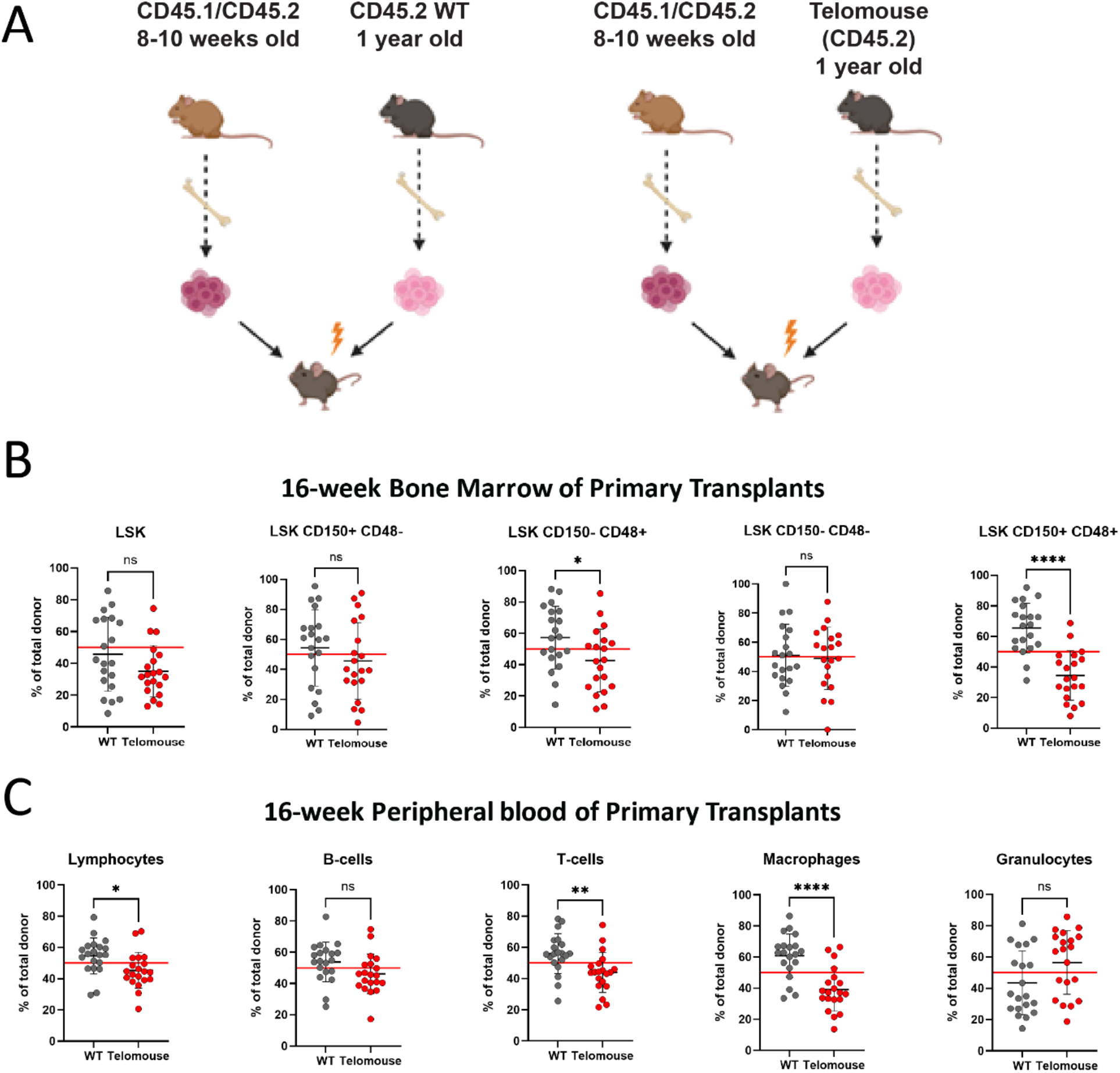
Aged Telomice show a reduction in both circulating immune cells and bone marrow stem cells in a bone marrow competition assay. (A) Schema of experimental design. Aged WT and Telomouse bone marrow cells were competed against young WT bone marrow cells. (B) Analysis of the hematopoietic stem cell compartment of recipient mice 16 weeks after bone marrow transplant. Stem and progenitor cells are identified as in Fig. 2, and percentages represent the proportion of WT and Telomouse donor cells with respect to the total proportion of donor cells (CD45.2 + CD45.1/CD45.2) within each population in the host mouse. (C) Composition of peripheral blood (PB) lymphocytes following 16 weeks of repopulation. Red lines indicate the percentage that would represent equal contribution by aged and young donors. Student’s T-test was used for statical analysis. N = 10 WT mice and 10 Telomice (F12). ****, p ≤ 0.0001; ***, p ≤ 0.001; **, p ≤ 0.01; *, p < 0.05.

Genome integrity and replicative health of a cell are determined by the shortest telomeres, as below a certain length the protective shelterin complex cannot suppress the DNA damage response at the DNA end, which limits further cell proliferation (Palm and de Lange 2008). We sought to determine if the loss of functional telomere length in some of a cell’s chromosomes accounted for the mechanism by which stressed Telomouse bone marrow cells failed to regenerate appropriately. We analyzed metaphase spreads on WT or Telomouse bone marrow cells. Fluorescent in situ hybridization (FISH) was performed to probe the spread chromosomes for the canonical telomeric G-strand repeats (TTAGGG). This allowed us to determine the number of chromosome ends without detectable telomeric signal (signal-free ends) in a cell, as well as visualize other genetic aberrations such as chromosome fusions. We found that even at homeostasis, Telomouse bone marrow cell chromosomes displayed a significant number of signal-free ends (Fig.5A, B), while the other chromosomal aberrations were not significant, consistent with the metaphase FISH results on Telomouse immortalized mouse embryonic fibroblasts (Smoom et al. 2023). FISH for telomere repeat arrays has been shown to detect telomeres below 1 kb in length (Aubert et al. 2012; Montpetit et al. 2014), indicating that the signal-free chromosome ends seen in Telomouse bone marrow cells represent extremely short telomeres. This suggests that while these cells may not be in crisis yet, they are already very close. Interestingly, no metaphase spreads could be generated from Telomouse bone marrow from 5-FU treated mice, which we suspect is due to cells being unable to successfully progress through mitotic checkpoints within the experimental timeframe (data not shown).

**Figure 5.**
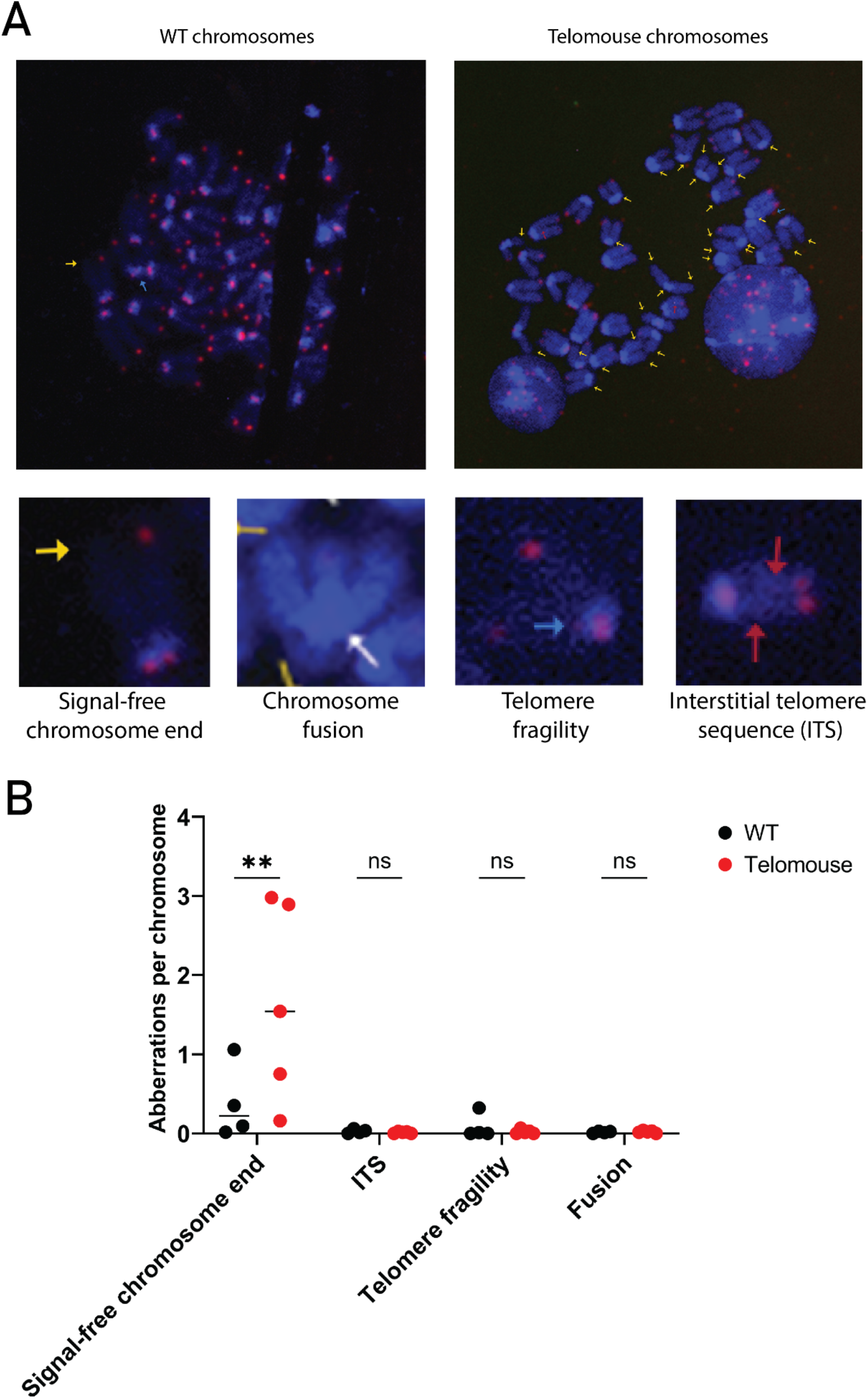
A subset of Telomouse bone marrow cells exhibit extremely short telomeres. (A) Representative images of metaphase chromosomes from wild type or Telomice (blue= DAPI, red= fluorescent *in situ* hybridization for telomere repeats). Chromosomal aberrations are indicated with colored arrows (yellow arrow = no telomeric signal at a chromosome arm end, white arrow = chromosomal fusion, blue arrow = telomere fragility, red arrow = interstitial telomere sequence [ITS]). (B) Quantification of chromosomal aberrations per chromosome from wild type (N= 4) or Telomice (N=5, F17). Between 1-3 fields of view were analyzed and averaged per mouse. Analysis was performed using a two-way ANOVA with Šídák’s multiple comparisons test. **, p ≤ 0.01

In conclusion, employing the recently established Telomouse model, which is a *M. musculus* C57BL/6 sub-strain with human-length telomere repeats, we addressed the extent to which this telomere size limits the regenerative capacity of the hematopoietic stem cell compartment. We note no significant impact under homeostatic conditions, where the proportions of the various hematopoietic stem cells and multipotent progenitor populations are apparently unchanged. Likewise, bone marrow stem cells from young Telomice perform nearly equivalent to wild type mice in a competitive repopulation assay. However, under conditions of high replicative stress, whether following 5-FU treatment, undergoing a secondary bone marrow transplant, or in aged mice, Telomice with human length telomeres are at a competitive disadvantage. We propose that the differences in the regenerative capacity of the hematopoietic stem cells system between humans and *M. musculus* can be explained in part by the difference in telomere lengths, and that the Telomouse is a better model for studying telomere dynamics than wild type mice.

## Materials and Methods

### Animals

Telomice were generated and maintained as described previously (Smoom et al. 2023). The Telomice analyzed here were between F11 and F17. Animals were maintained on 12-hour dark/light cycles with *ad libitum* access to standard rodent chow and water. The use of animals in this study followed the Guide for the Care and Use of Laboratory Animals, Laboratory Animal Ordinances, and the Animal Welfare Act. This study was approved by the University of Pennsylvania Institutional Animal Care and Use Committee.

### Bone marrow collection

Femurs and tibia were harvested from mice, their ends removed, and bone marrow cells extracted by spinning under submersion of 5% FBS/PBS (5% Fetal Bovine Serum in PBS). Cells were then filtered three times using a FACS tube with cell strainer cap (Falcon 352235) and resuspended in 5% FBS/PBS.

### Flow Cytometry

Peripheral blood analysis was performed at 8- and 15-16-weeks post-transplant. Approximately 75 μl of blood was collected from retroorbital venous sinus of the mouse, treated with ACK (Ammonium-Chloride-Potassium) Lysing Buffer (Lonza, BP10-548E) two times for 10 minutes, then washed and filtered using a 70 µm cell strainer (BD Biosciences). For analysis of 5-FU treated mice, bone marrow cells were isolated as above and a lineage depletion was performed by incubating cells in biotinylated B220, CD3e, CD11b, Gr1, Ter119 (eBiosciences) and streptavidin-microbeads (Miltenyi Biotech). Lineage-negative cells were run through MS columns on MACS magnetic separator (Miltenyi Biotech).

Cells were washed with 5% FBS/PBS and resuspended in 100 μl antibody cocktail (Supplementary Table S1) for 30 minutes at 4°C. Cells were then strained through a 70 µm filter and stained with 0.5 mg/mL DAPI in FBS. Flow cytometry was performed on a FACSAria instrument, and data were analyzed using Fowjo 9.0.1 (Tree Star, Inc., Ashland, OR).

### 5-FU treatment

Seven-to nine-week-old mice were injected intraperitoneally with suspended 5-fluorouracil (150 mg/kg body weight; Thermo Scientific cat # 228440010) every 7 days for three weeks.

### Bone marrow transplantation

B6.SJL-*Ptprc*^*a*^*Pepc*^*b*^/BoyJ (CD45.1) mice were administered 900 cGy of irradiation using GammaCell 40 ^137^Cs irradiator in a split dose separated by 3-4 hours. 2x10^6^ total bone marrow cells resuspended 0.2 mL 5% FBS/PBS were injected retro-orbitally while mice were under anesthesia. Mice were monitored for signs of dehydration throughout treatment and given subcutaneous injection of 0.2 ml 0.9% normal saline for relief. Mice were also maintained on antibiotic diet food containing sulfamethoxazole and trimethoprim (Lab Diet, p/n 5053) for two weeks post-transplantation. CD45.1/2 mice used as competitors were generated by crossing B6.SJL-*Ptprc*^*a*^*Pepc*^*b*^/BoyJ with C57BL/6J mice.

### Metaphase spreads/telomere FISH

Bone marrow cells were collected as outlined above and were plated in RPMI 1640 media supplemented with 10% FBS and 1% penicillin-streptomycin. Bone marrow cells were incubated for 2 hours before media was supplemented with Colcemid (0.1 μg/mL; Thermo Scientific cat # 15212012). Colcemid-treated cells were incubated for 30 minutes to allow dividing cells to arrest in metaphase, then cells were removed using trypsin and collected by gentle centrifugation (5 min, 100 g). Cell pellets were gradually resuspended in hypotonic media (0.075 M KCl) warmed to 37°C. Additional hypotonic media was added until the volume was at least ten times that of the cell pellet. Samples were then incubated for 25 minutes at 37°C to promote cell swelling. Following the incubation, samples were spun down and cell pellets were slowly resuspended in fixative (3:1 methanol: glacial acetic acid). Cells were washed twice and were then resuspended in fixative. These samples were then dropped from a height of 0.7 meters onto a glass slide that had been pre-treated with L-proline and steamed. Slides were dried at 65°C and subsequently stored at room temperature until hybridization.

To perform telomeric FISH hybridization, slides were first dehydrated using a graded ethanol series (70%, 95%, 100%) then air-dried. Sample areas were then covered with hybridization mix, which consisted of 14 μL deionized formamide (EMD Millipore, S4117), 2 μL blocking solution (Roche, REF# 11096176001, prepared according to manufacturer recommendations), 1 μL 1M Tris-HCl pH 8.0, 1 μL MgCl_2_ buffer (82 mM Na_2_HPO_4_, 9 mM citric acid, 25 mM MgCl_2_), 1 μL telomere probe (Panagene, F1002-5), and 1 μL molecular biology-grade water per slide. Slides were heated at 80°C for 10 minutes in the dark, then hybridized at room temperature in the dark overnight. After hybridization, slides were washed twice with a 70% formamide, 10 mM Tris-HCl pH 8.0 solution then thrice with a 50 mM Tris-HCl pH 8.0, 150 mM NaCl, 0.08% Tween-20 solution at room temperature. Slides were submerged in deionized water to remove excess salt and then dried in the dark at room temperature. Dry slides were coated with Vectashield + DAPI and a coverslip, followed by curing in the dark at room temperature overnight.

## Competing Interest Statement

The authors declare no competing interests.

## Acknowledgements

We thank Mark Tigue for help with the mouse experiments. This work was supported in part by the Israel Science Foundation grants 2071/18 and 1342/23 to Y.T., the NIDDK through 1R01DK139049-01 and R01-CA249929 to K.H.K., R01HL-091724-31 to N.A.S., and by the Israel-UK-Palestine GROWTH Fellowship to R.S.

## Author Contributions

M.R, J.T. and V.O. performed the mouse experiments. R.S. developed the protocol for FISH hybridization, which M.R. performed. M.R., J.T., R.S., N.A.S., Y.T. and K.H.K. wrote and edited the manuscript.

## Supplemental figures

**Figure S1.**
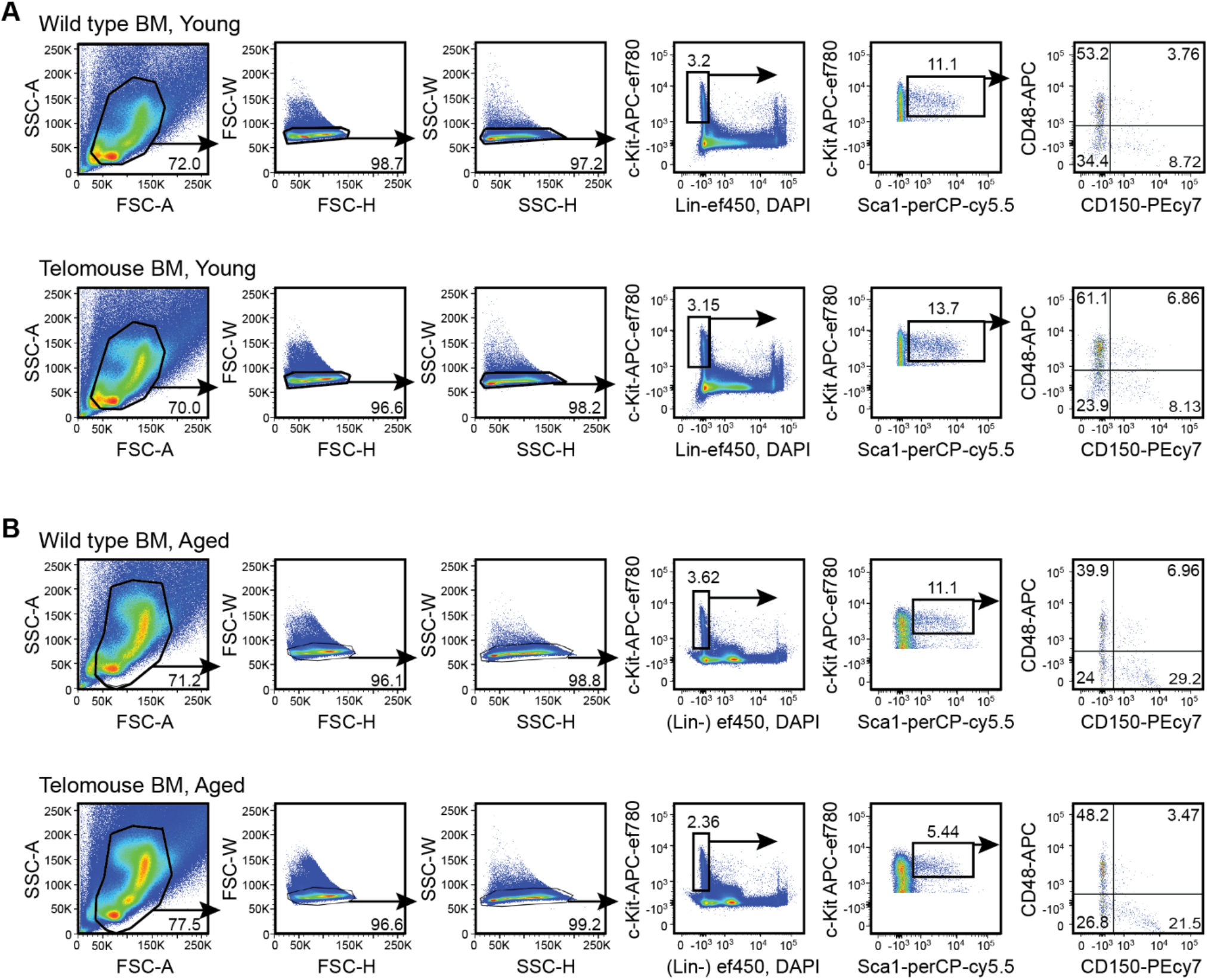
Flow cytometric analysis of hematopoietic stem and progenitor cells in the bone marrow of young (A) and aged (B) wild type and Telomice (RTEL^K/K^). Percentages are the portion of cells within each gate.

**Figure S2.**
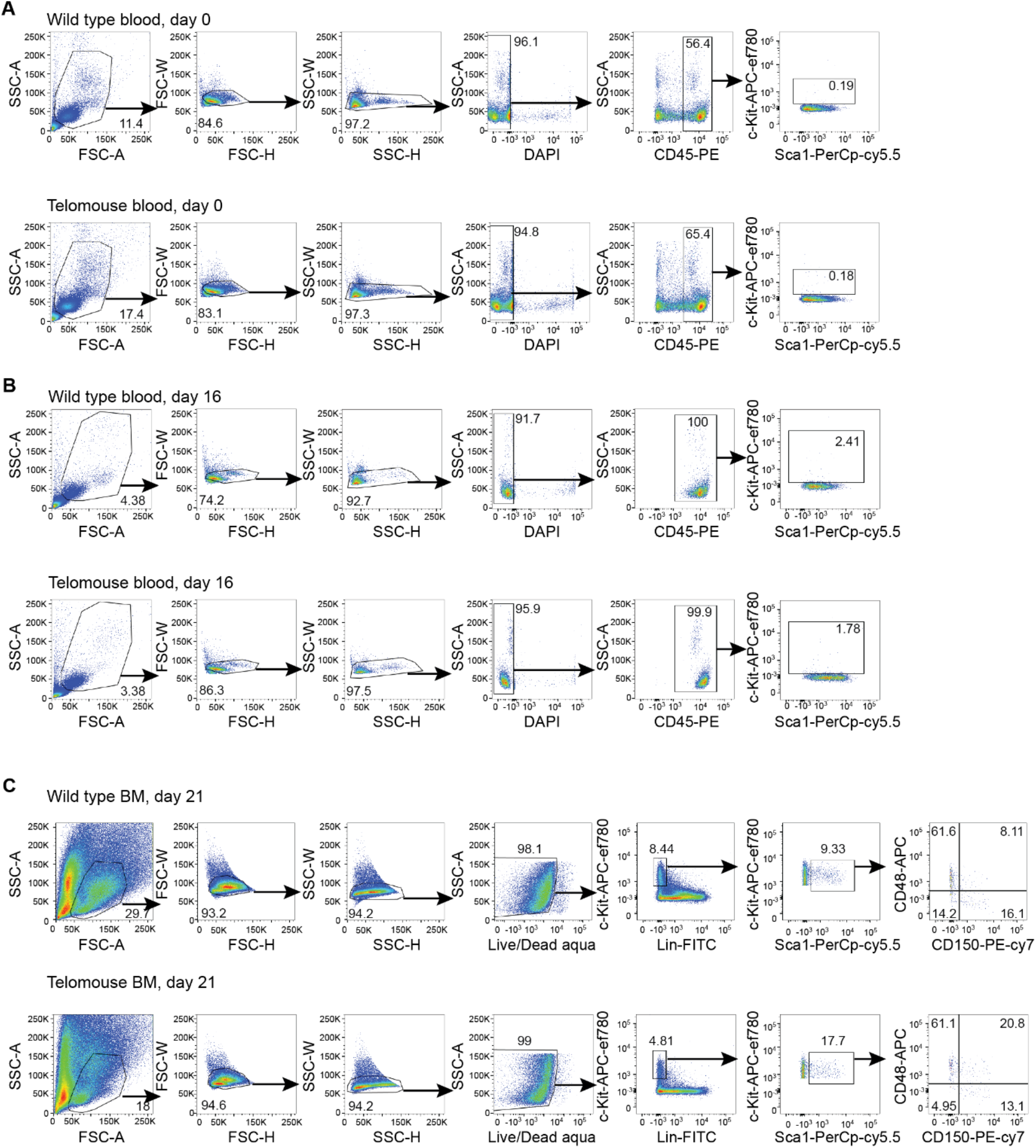
Flow cytometric analysis of blood and bone marrow for 5-FU experiments. A. Peripheral blood analysis at day 0 (untreated mice) showing the baseline percent of c-Kit+ cells. B. Peripheral blood analysis of c-Kit+ cells 16 days-post the initiation of 5-FU treatment. C. Bone marrow analysis of hematopoietic stem and progenitor cells 21 days-post the initiation of 5-FU treatment.

**Figure S3.**
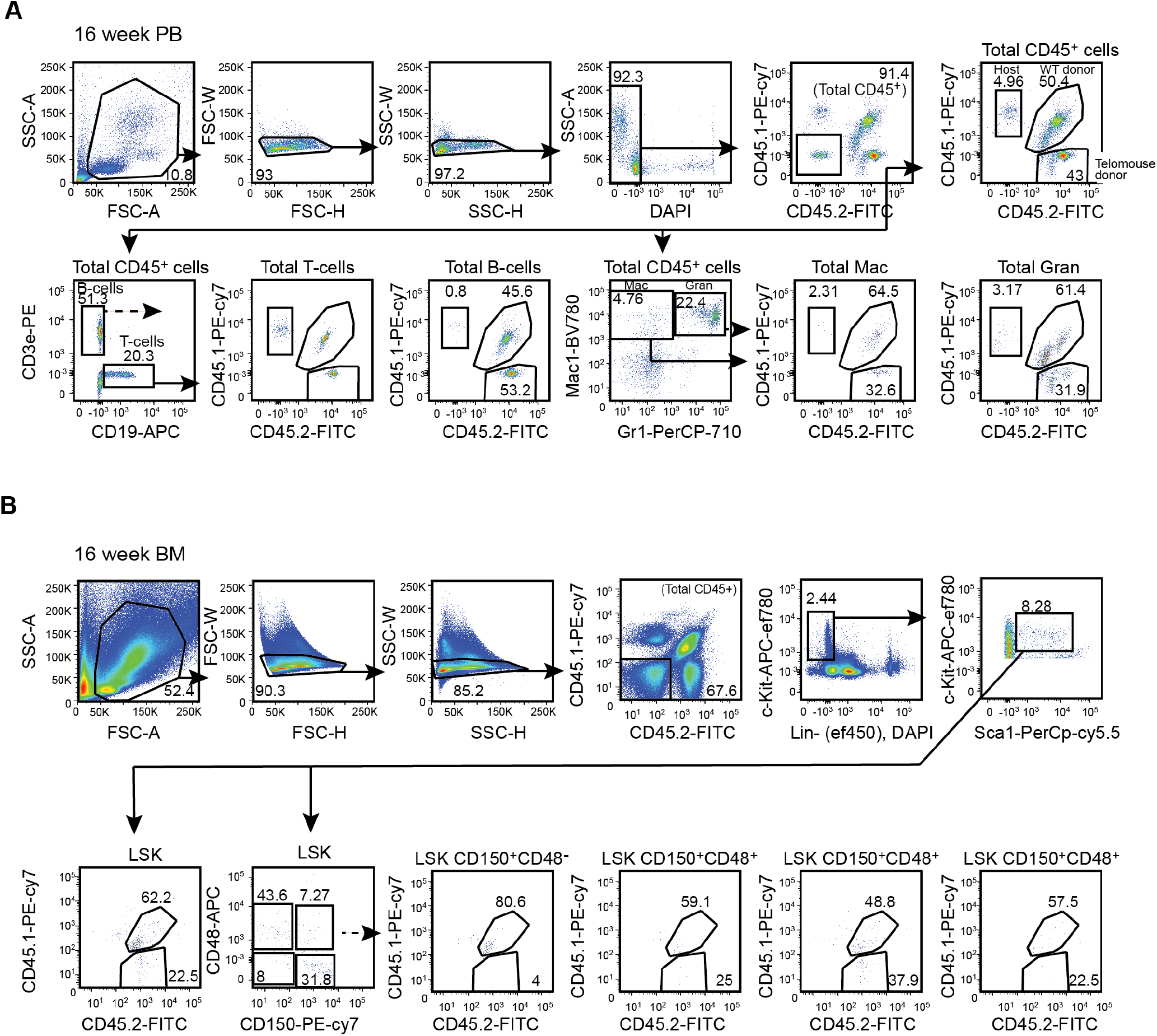
Gating strategy used to examine donor (Telomouse, CD45.2+) and competitor (wild type, CD45.1+/CD45.2+) cells 16-weeks post-transplant. A. Gating strategy for donor and competitor lineage+ in the peripheral blood. B. Gating strategy for donor and competitor hematopoietic stem and progenitor cells.

**Supplemental Table S1.** Antibodies used for flow cytometry

| antibody | ab registry | fluorochrome |
| --- | --- | --- |
| CD45.2 | AB_465062 | FITC |
| CD45.1 | AB_469629 | PE-cy7 |
| CD45.1 | AB_465675 | PE |
| Mac1 | AB_2722926 | SB780 |
| Mac1 | AB_1582236 | e450 |
| Gr1 | AB_1548797 | e450 |
| Gr1 | AB_2848343 | Percp-e710 |
| CD19 | AB_1659676 | APC |
| CD3e | AB_465497 | PE |
| CD3e | AB_10735092 | e450 |
| B220 | AB_1548761 | e450 |
| Ter119 | AB_1518808 | e450 |
| c-Kit | AB_1272177 | APC-e780 |
| Sca1 | AB_914372 | Percp-cy5.5 |
| Sca1 | AB_469669 | PE-cy7 |
| CD48 | AB_571997 | APC |
| CD150 | AB_439797 | PE-cy7 |
| IL7ra | AB_2016629 | e450 |
| CD34 | AB_10596826 | APC |
| CD16/32 | AB_1877268 | Percp-cy5.5 |

